# E2F1 methylation by SETD6 regulates SETD6 expression via positive feedback mechanism

**DOI:** 10.1101/2023.06.27.546651

**Authors:** Margarita Kublanovsky, Gizem T. Ulu, Sara Weirich, Nurit Levy, Michal Feldman, Albert Jeltsch, Dan Levy

## Abstract

The protein lysine methyltransferase SETD6 has been shown to influence different cellular activities and are critically involved in the regulation of diverse developmental and pathological processes. However, the upstream signal which regulates the mRNA expression of SETD6 is not known. Bioinformatics analysis revealed that SETD6 promoter has a binding site for the transcription factor E2F1. Using various experimental systems, we confirmed that E2F1 binds to the SETD6 promoter and regulates SETD6 mRNA expression. Our further observation that this phenomenon is SETD6 dependent, suggested that SETD6 and E2F1 are linked. We next demonstrate that SETD6 mono-methylates E2F1 specifically at K117 *in-vitro* and in cells. Finally, we show that E2F1 methylation at K117 positively regulates the expression level of SETD6 mRNA. Depletion of SETD6 or overexpression of E2F1 K117R mutant which canot be methylated by SETD6, reverses the effect. Taken together, our data provide evidence for a positive feedback mechanism which regulates the expression of SETD6 by E2F1 in a SETD6 methylation dependent manner and highlight the importance of protein lysine methyltransferases and lysine methylation signaling in the regulation of gene transcription.

## Introduction

Protein lysine methyltransferases (PKMTs) catalyze lysine methylation, an increasingly important post-translational modification that regulates various signaling pathways (1,2). All PKMTs use S-adenosyl-L-methionine (SAM) as a methyl donor to methylate their target, the ε-amino group of lysine resulting in a mono-, di- or trimethylated lysine (Kme1, Kme2 or Kme3) (3,4). SET domain-containing protein 6 (SETD6) is a 53 kDa PKMT containing a catalytic SET domain and a Rubisco substrate-binding domain (5,6) which is encoded on chromosome 16 (16q21). SETD6 was originally identified as a mono-methyltransferase that directly methylates RelA (p65), a subunit of the nuclear factor κB (NF-κB) complex. This methylation was shown to suppress the activation of NF-κB target genes (7). Since then, SETD6 has been implicated in various biological processes such as gene expression regulation, chromatin remodeling, and cell cycle progression (5,7–19). SETD6 has also been linked to several developmental steps and pathological conditions. For example, SETD6 was shown to be essential for memory consolidation, regulation of gene expression patterns, and orderly spine morphology in the rat hippocampus (20). SETD6-mediated mono-methylation of BRD4 at K99 (10) was demonstrated to regulate human papillomavirus (HPV) transcription, genome replication, and segregation by binding of BRD4 to the E2 protein (21). In diabetic nephropathy, a chronic complication of diabetes, down-regulation of SETD6 protected the cells from apoptosis and mitochondrial dysfunction (21). Furthermore, SETD6 function has been associated with several cancer types (8,22,23). In bladder cancer, SETD6 is upregulated and promotes cell survival through the NF-κB pathway (23). In contrast, SETD6-mediated methylation of PAK4 inhibited cell migration and invasion in breast cancer (17). Due to its association with tumorigenic hallmarks, SETD6 may serve as an attractive target for therapeutic intervention (8,22,23).

The E2F family of transcription factors is an important downstream effector of the retinoblastoma tumor-suppressor gene product (pRB) and plays a crucial role in regulating cell-cycle progression. Additionally, E2Fs participate in a wide range of biological processes, such as differentiation, mitosis and the mitotic checkpoint, DNA replication, DNA-damage checkpoints, DNA repair, and apoptosis (24–26). The E2F family consists of eight members, namely E2F1-E2F8, which share the highest degree of homology in their DNA binding domain explaining their ability to bind to a unique E2F consensus sequence (27). However, experimental evidence suggests that different members of the E2F family regulate distinct yet overlapping sets of target genes (28–30). The specificity of E2F binding to an individual binding site can be influenced by the DNA sequence as well as interactions with other transcription factors that are bound to adjacent regulatory elements (27,31).

The observation that E2F1 is closely involved in regulating cellular processes which are also controlled by SETD6, suggested a potential functional cross-talk between SETD6 and E2F1. Here we demonstrate that SETD6 mono-methylates E2F1 at K117 *in-vitro* and in cells. We further show that E2F1 methylation increases the occupancy of E2F1 at the SETD6 locus, resulting in the increased expression of SETD6 mRNA. Together, our findings suggest a new mechanistic dimension for the selective regulation of SETD6 mRNA expression which is mediated by E2F1 activity via SETD6-dependent E2F1 methylation in a positive feedback mechanism.

## Materials and methods

### Plasmids

E2F1 sequence was amplified by PCR from a plasmid provided by Prof. Assaf Rudich (BGU) and subcloned into pcDNA3.1 3xFlag plasmid using primers indicated in table 1. For viral infections, E2F1 was cloned into the pWZL-Flag plasmid (7). For recombinant protein purification, E2F1 was cloned into pET-SUMO and pGEX-6p1 plasmids (7). To generate E2F1 mutants, site-directed mutagenesis on an E2F1 wild-type vector was performed using primers indicated in table 1, followed by DNA sequencing for confirmation. All E2F1 lysine mutants were cloned into pET-SUMO, pGEX-6p1, pcDNA3.1 3xFlag and pWZL-Flag plasmids. SETD6 promoter sequence was amplified by PCR using primers indicated in table 1, and subcloned into pGL3 plasmid.

**Table 1.**
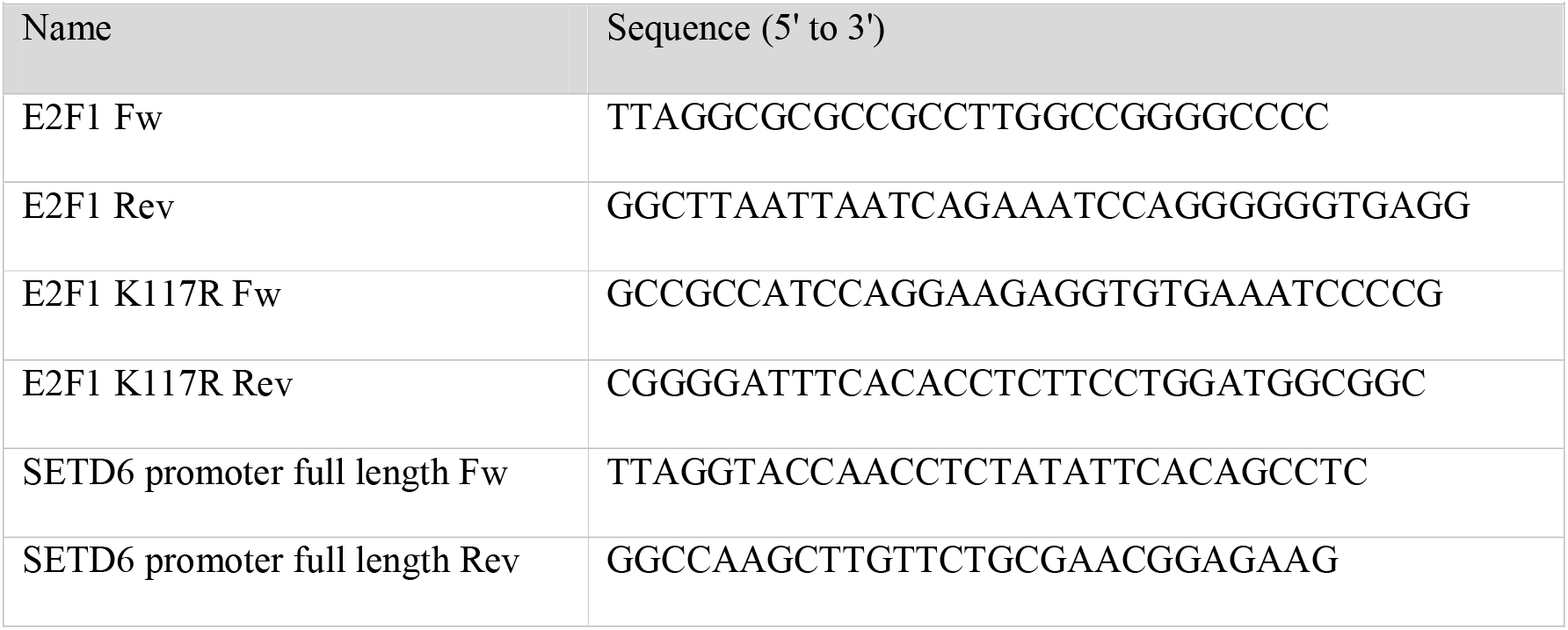
Primers for cloning and mutagenesis.

### Cell lines, Transfections, Infections and Treatments

HEK293 and DU145 (purchased from ATCC, kindly provided by Prof. Etta Livneh, BGU) cells were maintained in Dulbecco’s modified Eagle’s medium (Sigma, D5671) with 10% fetal bovine serum (FBS) (Gibco), penicillin-streptomycin (Sigma, P0781), 2 mM L-glutamine (Sigma, G7513) and non-essential amino acids (Sigma, M7145), at 37°C in a humidified incubator with 5% CO2 as previously described (32). Cell transfection was performed using Mirus reagents (TransIT-LT1 or TransIT-X2), according to the manufacturer’s instructions. The mCherry tagged SETD6 WT or Y285A and EGFP-tagged E2F1 WT or K117R were co-transfected into cells using polyethyleneimine (Thermo Fisher Scientific) according to manufacturer’s instructions. For the DU145 CRISPR/Cas9 SETD6 knock-out, two different gRNAs for SETD6 (Table 2) were cloned into lentiCRISPR plasmid (Addgene, #49535). Following transduction and puromycin selection (2.5 µg/ml), single clones were isolated, expanded and validated by sequencing.

**Table 2.**
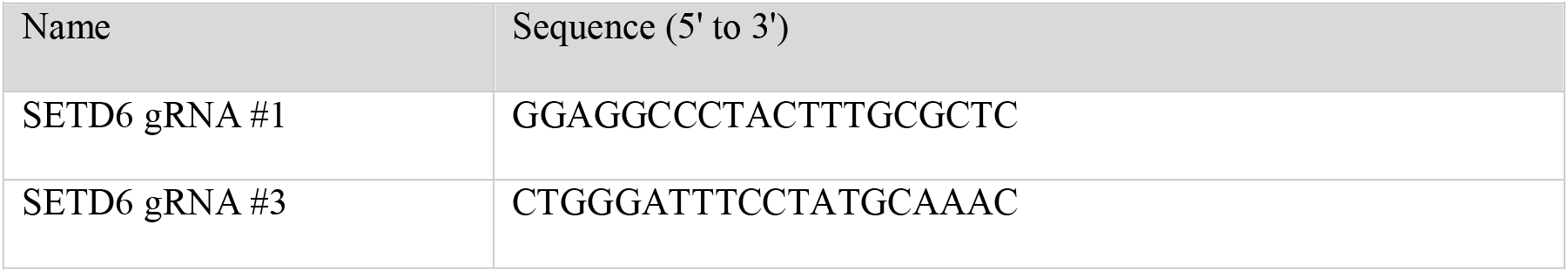
gRNAs for CRISPR-Cas9 KO cells.

### Recombinant Proteins

*Escherichia coli* Rosetta strain (Novagen) was transformed with a plasmid expressing His-SUMO-/GST-E2F1 WT or K117R, His-SETD6 WT or Y285A mutant proteins, and the transformed cells were grown in LB medium. Bacteria were harvested by centrifugation after IPTG induction. The bacteria overexpressing His-tagged proteins were re-suspended in lysis buffer containing 10 mM Imidazole, 1 mM PMSF and 0.1 v/v % Triton X-100 in phosphate-buffered saline (PBS), incubated with 0.25 mg/ml lysozyme for 30 minutes on ice, and followed with lysis by sonication on ice (25% amplitude, 1 min total, 10/5 sec ON/OFF). His-tagged proteins were purified using Ni-NTA beads (Pierce) or on a HisTrap column (GE) with the ÄKTA gel filtration system using PBS as wash buffer. Proteins were eluted by 0.5 M imidazole in PBS followed by dialysis to 10% glycerol in PBS. The bacteria overexpressing GST-E2F1 were re-suspended in lysis buffer containing 50 mM NaCl, 5 mM EDTA, 0.15 mM PMSF and 1% Triton X-100 in 50 mM Tris-HCl pH 8. GST-E2F1 was purified on glutathione-sepharose 4B (GE) or on a GSTrap column (GE) with the ÄKTA gel filtration system using PBS as wash buffer. Proteins were eluted with 10 mg/ml reduced glutathione (Sigma) in 50mM Tris pH 8. Recombinant GST SETD6 was expressed and purified as previously described (7).

### Antibodies, Western Blot Analysis and Immunoprecipitation

Primary antibodies used were anti-Flag (Sigma, F1804), anti-Actin (Abcam, ab3280), anti-Pan-Kme1 (Cell signaling, 14679), anti-GST (Abcam, ab9085), anti-His (ThermoFisher scientific, rd230540a), anti-SETD6 (Genetex, GTX629891), anti-E2F1 (SantaCruz, KH95), anti GFP antibody (Clontech, lot. 1404005) and anti-H3 (Abcam, ab10799). HRP-conjugated secondary antibodies, goat anti-rabbit, goat anti-mouse, and streptavidin-HRP were purchased from Jackson ImmunoResearch (111-035-144, 115-035-062 respectively). For Western blot analysis, cells were homogenized and lysed in RIPA buffer (50 mM Tris-HCl pH 8, 150 mM NaCl, 1% Nonidet P-40, 0.5% sodium deoxycholate, 0.1% SDS, 1 mM DTT, and 1:100 protease inhibitor mixture (Sigma)). Samples were resolved on 8-12% SDS-PAGE, followed by Western blot analysis.

The polyclonal E2F1-K117me1 antibody was generated by Abmart Inc. (Shanghai, China) using a GRHPGKme1GVK epitope identification peptide. For validation of its specificity, SPOT arrays were blocked in 5% milk in 1X TBS-T solution for 1 hour. Then, the array was incubated with the primary E2F1-K117me1 antibody solution (1:2000) overnight at 4 °C. The next day, the array was washed 3 times for 5 minutes with 1X TBS-T solution incubated with the secondary antibody solution anti-rabbit HRP (Na934v, GE Healthcare; 1:5000) for 1 hour at room temperature. After washing again, the signal was detected by chemiluminescence after the addition of Pierce™ ECL Western Blotting substrate.

For immunoprecipitation, proteins extracted from cells were incubated overnight at 4°C with FLAG-M2 beads (Sigma, A2220) or with antibody of interest, to which Magna ChIP™ Protein A+G Magnetic Beads (Millipore, 16-663) were added for 2hour at 4°C. The beads were then washed once with PBS and submitted to SDS-PAGE and Western blot analysis.

### GFP-Trap assay

HEK293 cells were grown in Dulbecco’s modified Eagle’s medium (Gibco) supplemented with 10% fetal bovine serum, 1% penicillin/streptomycin, and 5% L-glutamine (Sigma-Aldrich). The mCherry tagged SETD6 WT or Y285A and EGFP-tagged E2F1 WT or K117R were co-transfected into HEK293 cells using polyethyleneimine (Thermo Fisher Scientific) according to manufacturer’s instructions. 72 hours after transfection, cells were washed with 1X PBS buffer (Gibco) and harvested by centrifugation at 500g for 5 min. For methylation analysis, the GFP-fused substrate proteins were immunoprecipitated from the cell extract using GFP-Trap A beads (Chromotek) according to the manufacturer’s instructions. The eluted protein samples were separated by 16% SDS-PAGE and Western Blot analysis was performed using the primary E2F1 K117me1 antibody solution (1:2000) and the secondary anti-rabbit HRP solution (1:5000). Equal loading of the GFP tagged E2F1 proteins was determined by the GFP antibody (Clontech, lot. 1404005; 1:2000) the secondary antibody anti-rabbit HRP (Na934v, GE Healthcare; 1:5000).

### Synthesis of Peptide SPOT Arrays

Peptide arrays were generated by the SPOT synthesis method using the Autospot Multipep peptide array synthesizer (Intavis AG). Each peptide spot, with a diameter of 2 mm, contained approximately 9 nmol of peptide (Autospot Reference Handbook; Intavis AG). The successful synthesis of peptide arrays was verified by bromphenol blue staining. Each spot comprises 15-amino acid long peptides with different residues surrounding the potential methylation target site.

### Methylation of peptide SPOT arrays

Peptide SPOT arrays were preincubated in methylation buffer containing 20 mM Tris/HCl pH 9 and 5 mM DTT for 5 min on a shaker. Afterwards, the SPOT arrays were incubated in methylation buffer containing additionally 50 nM SETD6 and 0.76 µM labelled [methyl-^3^H]-AdoMet (Perkin Elmer Inc.) for 1 hour at room temperature. Next, the arrays were washed five times for 5 minutes with 100 mM NH_4_HCO_3_ and 1% SDS. After washing, the arrays were incubated in Amplify NAMP100V solution (GE Healthcare) for 5 minutes. Then, the arrays were exposed to HyperfilmTM high-performance autoradiography films (GE Healthcare) in the dark at −80 °C for different exposure times and developed using an Optimus TR developing machine.

### *In-vitro* protein methylation Assay

1.6 µM of GST-tagged E2F1 WT or mutant was incubated with 0.2 μM of His-SUMO-tagged SETD6 WT or mutant in methylation buffer (20 mM Tris/HCl pH 9 and 5 mM DTT), supplemented with 0.76 µM labelled [methyl-^3^H] -AdoMet (Perkin Elmer) for 3 hours at 25°C. The reaction was stopped by the addition of SDS-PAGE loading buffer and heating for 5 min at 95 °C. Afterwards, the samples were separated by 16% SDS-PAGE followed by the incubation of the gel in amplify NAMP100V (GE Healthcare) for 1 h on a shaker. In the next step, the gel was dried for 2 hours at 70 °C under vacuum. The signals of the transferred radioactively labelled methyl groups were detected by autoradiography using a Hyperfilm^TM^ high performance autoradiography film (GE Healthcare) at −80 °C in the dark. The film was developed with an Optimax Typ TR machine after different exposure times.

For the non-radioactive methylation assay, the reactions were supplemented with 1 mM of non-radioactive AdoMet (Sigma-Aldrich). The reaction was stopped by the addition of SDS-PAGE loading buffer and heated for 5 min at 95 °C. Afterwards, the samples were separated by 16% SDS-PAGE followed by Western Blot analysis.

### Enzyme-Linked Immunosorbent Assay (ELISA)

His-SUMO-E2F1 (2 µg), MBP-RelA (2 µg) or BSA diluted in PBS were added to a 96-well plate (Greiner Microlon) and incubated for 1 hour at room temperature followed by blocking with 3% BSA for 30 min. Then, the plate was covered with 0.5 µg GST-SETD6, or GST protein (negative control) diluted in 1% BSA in Phosphate-buffered saline and Tween 20 (PBST) for 1 hour at room temperature. Plates were then washed and incubated with primary antibody (anti-GST, 1:4000 dilution) followed by incubation with HRP-conjugated secondary antibody (goat anti-rabbit, 1:2000 dilution) for 1 hour. Finally, TMB reagent followed by 1 N H_2_SO_4_ (stop solution) were added; the absorbance at 450nm was detected using Tecan Infinite M200 plate reader.

### RNA Extraction and Real-Time qPCR

Total RNA was extracted using the NucleoSpin RNA Kit (Macherey-Nagel). Then, 200 ng of the extracted RNA was reverse transcribed to cDNA using the iScript cDNA Synthesis Kit (Bio-Rad) according to the manufacturer’s instructions. The real-time qPCR primers were designed using the universal probe library assay design center (Roche) and UCSC Genome Bioinformatics (table 3). qPCR was performed using SYBR Green I Master (Roche) in a LightCycler 480 System (Roche) in a 384-well plate using the following cycling conditions: 5 min at 95°C, 45 cycles of amplification; 10 sec at 95°C, 10 sec at 60°C and 10 sec at 72°C, followed by melting curve acquisition; 5 sec at 95°C, 1 min at 65°C and monitoring up to 97°C, and finally cooling for 30 sec at 40°C. All samples were amplified in 4 or 5 replicates. Gene expression levels were normalized relative to GAPDH gene and controls of the experiment.

**Table 3.**
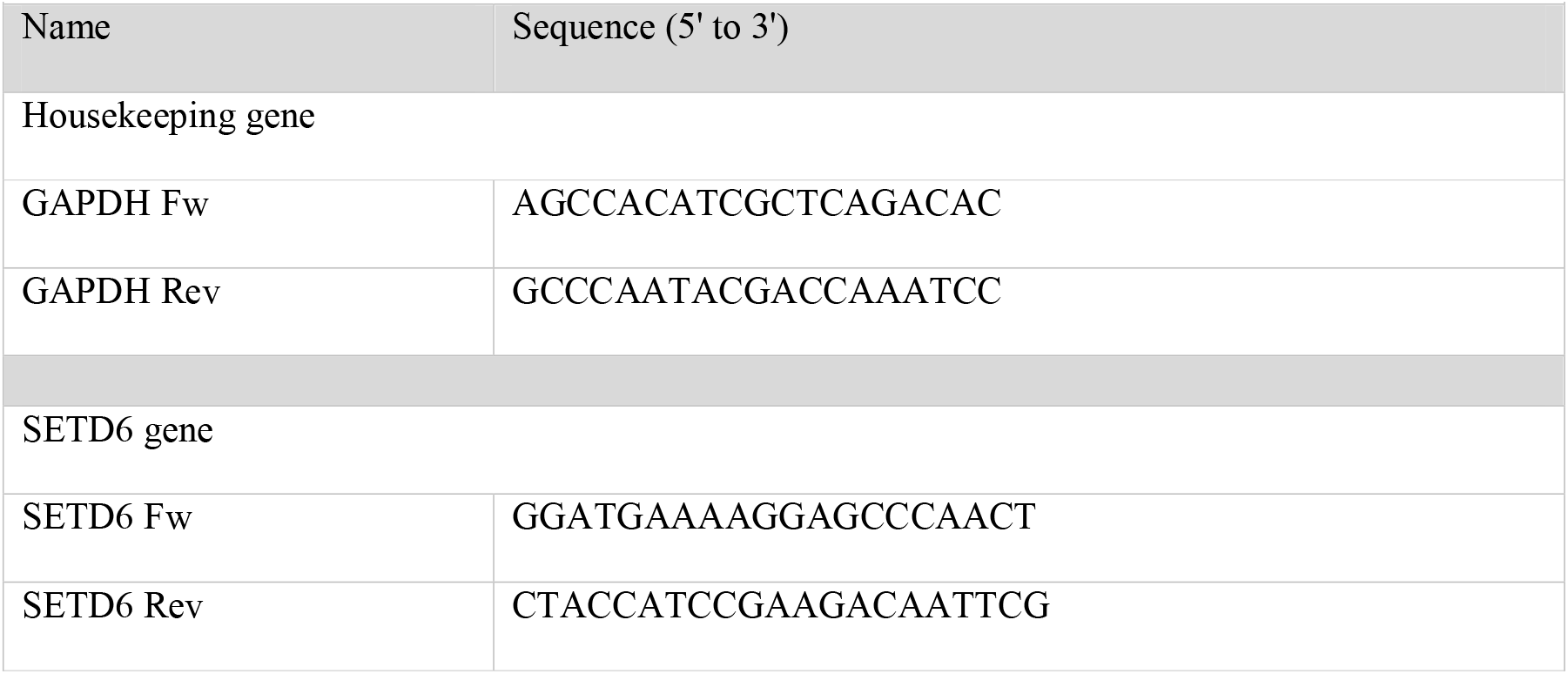
Primers for qPCR.

### Chromatin Extraction

Cells were cross-linked using 1% formaldehyde (Sigma) added directly to the medium and incubated on a shaking platform for 10 min at room temperature. The cross-linking reaction was stopped by adding 0.125 M glycine for 5 min. Cells were harvested and washed twice with PBS and then lysed in 0.5 ml cell lysis buffer (10 mM HEPES pH 7.9, 10 mM KCl, 1.5 mM MgCl_2_, 340 mM sucrose, 10% glycerol, 0.1% Triton X-100, 100 nM PMSF, 1 mM DTT, 1:200 protease inhibitor) for 8 min on ice. Samples were centrifuged (2,000g, 5 min, 4°C) and the pellets were washed with the buffer above, without protease inhibitor and centrifuged again. The nuclei pellets were lysed in 0.5 ml nuclei lysis buffer (3 mM EDTA, 0.2 mM EGTA, 1 mM DTT, 1:200 protease inhibitor) for 30 min on ice. Samples were centrifuged (2,000g, 5 min, 4°C) and the chromatin pellets were solubilized in 200 µl resuspension buffer (10 mM HEPES pH 7.9, 10 mM KCl, 1.5 mM MgCl_2_, 340 mM sucrose, 10% glycerol, 1:200 protease inhibitor, 1:200 benzonase nuclease enzyme (Sigma)) for 15 min at 37°C.

For protein-protein interaction analysis, the soluble chromatin was precleared with Magna ChIP™ Protein A+G Magnetic Beads (Millipore, 16-663) for 1 hour and then incubated overnight at 4°C with magnetic FLAG-M2 beads or the indicated antibody, then A+G magnetic beads were added for 2 hours at 4°C. The immunoprecipitated complexes were washed once with PBS. Immunoprecipitated complexes in protein sample buffer were resolved in SDS-PAGE and analyzed by Western blot.

### Chromatin Preparation and ChIP-qPCR

Cells were cross-linked using 1% formaldehyde (Sigma) added directly to the medium and incubated on a shaking platform for 10 min at room temperature. The cross-linking reaction was stopped by adding 0.125 M glycine for 5 min. Cells were harvested and washed twice with PBS and then lysed in 1 ml cell lysis buffer (20 mM Tris-HCl pH 8, 85 mM KCl, 0.5% Nonidet P-40, 1:100 protease inhibitor cocktail) for 10 min on ice. Nuclear pellets were resuspended in 200 μl nuclei lysis buffer (50 mM Tris-HCl pH 8, 10 mM EDTA, 1% SDS, 1:100 protease inhibitor cocktail) for 10 min on ice, and then sonicated (Bioruptor, Diagenode) at high power settings for 6 cycles, 6 min each (30 sec ON/OFF). Samples were centrifuged (20,000g, 15 min, 4°C) and the soluble chromatin fraction was collected. The chromatin fraction was diluted 5× in dilution buffer (20 mM tris-HCl pH 8, 2 mM EDTA, 150 mM NaCl, 1.84% Triton X-100, and 0.2% SDS). The chromatin fraction was precleared overnight at 4°C with A+G magnetic beads. The precleared sample was then immunoprecipitated with magnetic FLAG-M2 beads or A+G magnetic beads pre-conjugated with the indicated antibody. The immunoprecipitated complexes were washed according to the chromatin extraction protocol detailed above. DNA was eluted with elution buffer (50 mM NaHCO_3_, 140 mM NaCl, and 1% SDS) containing ribonuclease A (0.2 μg/μl) and proteinase K (0.2 μg/μl). Lastly, the DNA eluates were de-cross-linked at 65°C overnight with shaking at 900rpm and purified by NucleoSpin Gel and PCR Clean-up kit (Macherey-Nagel), according to the manufacturer’s instructions.

Purified DNA was subjected to qPCR using specific primers (Table 2). Primers were designed based on E2F1 occupancy (in correlation with H3K4me3, H3K27Ac, ATAC-seq and TF clusters in *SETD6* promoter locus) found in different ChIP-seq data previously published in NCBI GEO datasets by Ramos-Montoya *et al*. (33) (GEO accession: GSM1207898), Barfeld *et al*. (34) (GEO accessions: GSM1907203, GSM1907213), Bert *et al*. (35) (GEO accession: GSM947524) and Liu *et al*. (36) (GEO accession: GSM2186480) and viewed using Integrated Genomics Viewer software (IGV) (37). qPCR was performed using SYBR Green I Master (Roche) in a LightCycler 480 System (Roche). All samples were amplified in 4 replicates in a 384-well plate using the following cycling conditions: 5 min at 95°C, 45 cycles of amplification; 10 sec at 95°C, 10 sec at 60°C and 10 sec at 72°C, followed by melting curve acquisition; 5 sec at 95°C, 1 min at 65°C and monitoring up to 97°C, and finally cooling for 30 sec at 40°C. The results were normalized to input DNA and presented as % input. Primers used for the ChIP-qPCR are listed below in table 4.

**Table 4:**
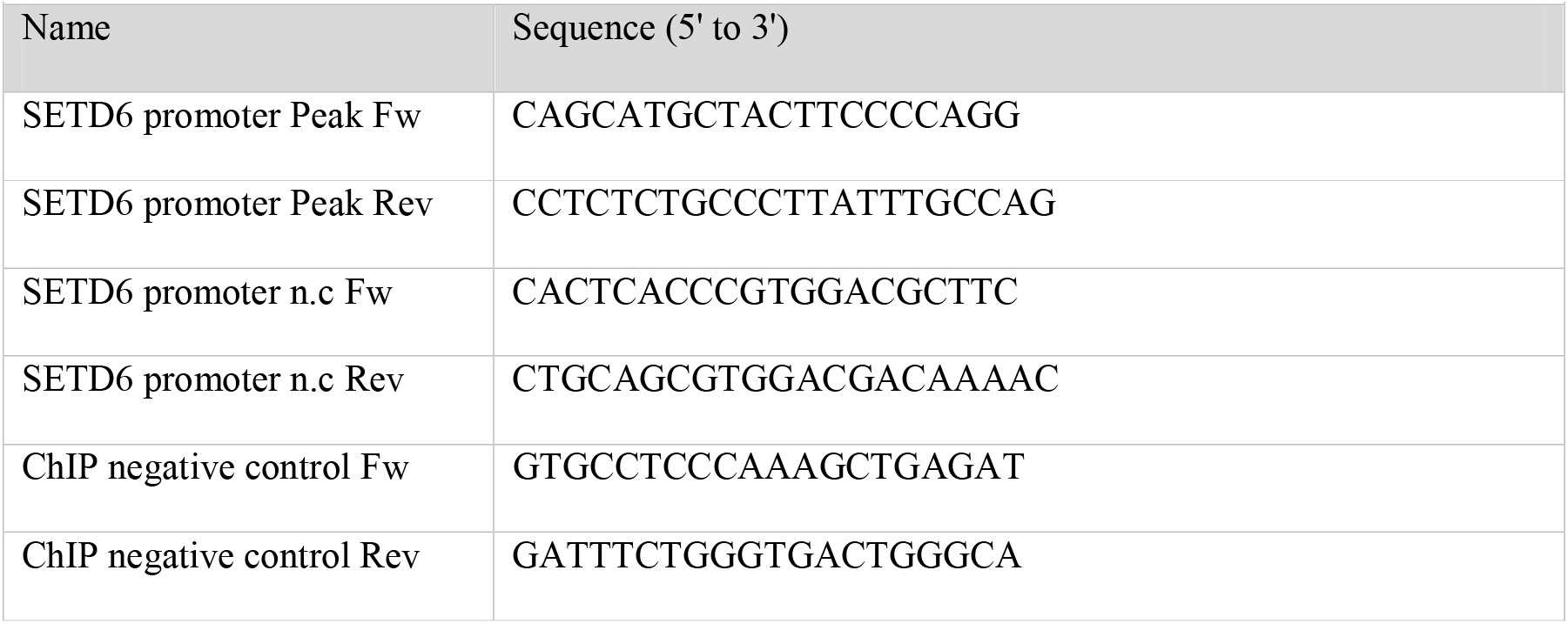
Primers for ChIP-qPCR.

### Dual-Luciferase Assay

DU145 cells were seeded in 24-well plates and transiently transfected with 0.1 µg Flag-E2F1 WT or K117R mutant or 25 mM siRNA, 0.1 µg firefly luciferase plasmid, containing SETD6 promoter variants, and 0.1 µg *Renilla* luciferase plasmid. Total amount of transfected DNA in each dish was kept constant by the addition of empty vector, as necessary. Cell extracts were prepared 30h after transfection, and firefly luciferase activity was measured with the Dual-Glo Luciferase Assay system (Promega) and normalized to that of *Renilla* luciferase.

### Statistical Analyses

Statistical analyses for all assays were analyzed with GraphPad Prism software, using Student’s two-tailed t-test (unpaired) or one-way analysis of variance (ANOVA) with a Tukey’s post hoc test.

## Results

### E2F1 regulates SETD6 mRNA expression

To investigate the potential mechanisms of *SETD6* gene regulation, we evaluated the presence of potential transcription factor binding sites at the *SETD6* promoter region (https://jaspar.genereg.net/) (38). We identified 48 TFs which potentially bind at *SETD6* promoter (Figure 1A, Supplementary table 1). Exploration of the Human Protein Atlas resource (https://www.proteinatlas.org/) summary on the pathology of SETD6 using immunohistochemical analysis, showed that malignant prostate cells had the highest rate of SETD6 protein expression, compared to other cancerous tissues (Supplementary Figure 1). To validate if these transcription factors are enriched in prostate cancer cells, we next searched publicly available ChIP-seq databases (Gene Expression Omnibus (GEO) DataSets https://www.ncbi.nlm.nih.gov/gds, Cistrome Data Browser http://cistrome.org/db/#/,) for selected TFs. Interestingly, our validations revealed selective enrichment of E2F1, ELK4 and MYC in prostate cells (Figure 1B). While we could not identify any enrichment for Myc and ELK4, we observed that E2F1 is highly enriched at the promoter area of *SETD6* which is in an open chromatin state marked by H3K27Ac and H3K4me3 (Figure 1B). We therefore decided to focus on the role of E2F1 in the expression regulation of SETD6 in prostate cells. Overexpression of Flag-E2F1 in DU145 cells followed by qPCR analysis revealed an increase of SETD6 mRNA expression levels (Figure 1C). We confirmed these results by qPCR analysis subsequent to treatment of DU145 cells with siRNA targeting endogenous E2F1 (Figure 1D). These results suggest that E2F1 stimulates SETD6 transcription.

**Figure 1.**
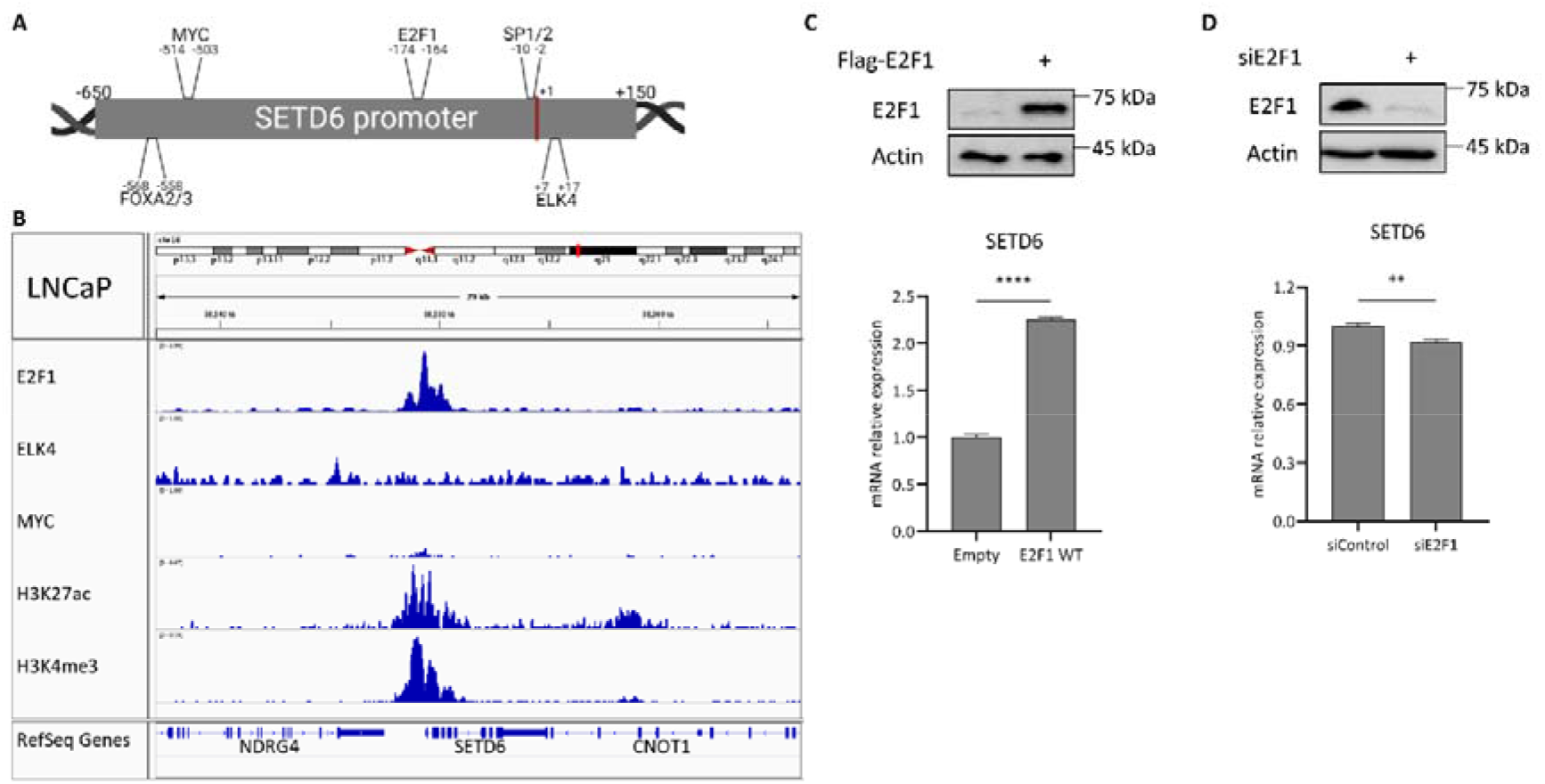
E2F1 is predicted to bind at SETD6 promoter and regulates SETD6 mRNA levels. **(A)** Scheme of SETD6 promoter and several human transcription factor binding sites that were predicted at the genomic region of the SETD6 promoter in the JASPAR database (https://jaspar.genereg.net/) with a relative profile score threshold of at least 90%. **(B)** SETD6 promoter genomic region (blue highlight) with E2F1 [GSM1656410], ELK4 [GSM1424528], MYC [GSM1907203], H3K27ac [GSM1907213], H3K4me3 [GSM1907211] ChIP-seq data in prostate cancer cell lines. Data were extracted by the Cistrome Data Browser: (http://cistrome.org/db/#/) and visualized using the IGV software. **(C, D)** DU145 cells were transfected with Flag-E2F1 WT **(C)** or with siRNA specifically targeting endogenous E2F1 **(D)**. 24 hours post-transfection, the SETD6 protein levels were assessed by WB (top) using the indicated antibodies, and the SETD6 mRNA expression levels were measured using RT-qPCR (bottom). mRNA expression levels were normalized to mRNA expression levels of GAPDH housekeeping gene. Error bars are s.e.m. Statistical analysis is based on 5 experimental repeats. **p≤0.002, ****p≤0.00001

### E2F1 activates SETD6 promoter transcription in a SETD6 dependent manner

To examine the possibility that E2F1 directly regulates *SETD6* transcription, we cloned the full-length promoter of SETD6 upstream to a luciferase reporter gene. Knock down of E2F1 expression using siRNA in DU145 cells, followed by luciferase assay confirmed E2F1 regulation of *SETD6* promoter activation (Figure 2A). In a reciprocal experiment, we found a significant elevation in the promotor activity after over-expression of Flag-E2F1 (Figure 2B). Interestingly, activation of luciferase transcription was lost in SETD6 KO cells, even when accompanied with E2F1 overexpression (Figure 2B). Sequence validation of the gRNAs is shown in Figure Supplementary Figure 2. Collectively, these data suggest that E2F1-mediated activation of SETD6 transcription is potentially regulated in a SETD6 dependent manner.

**Figure 2.**
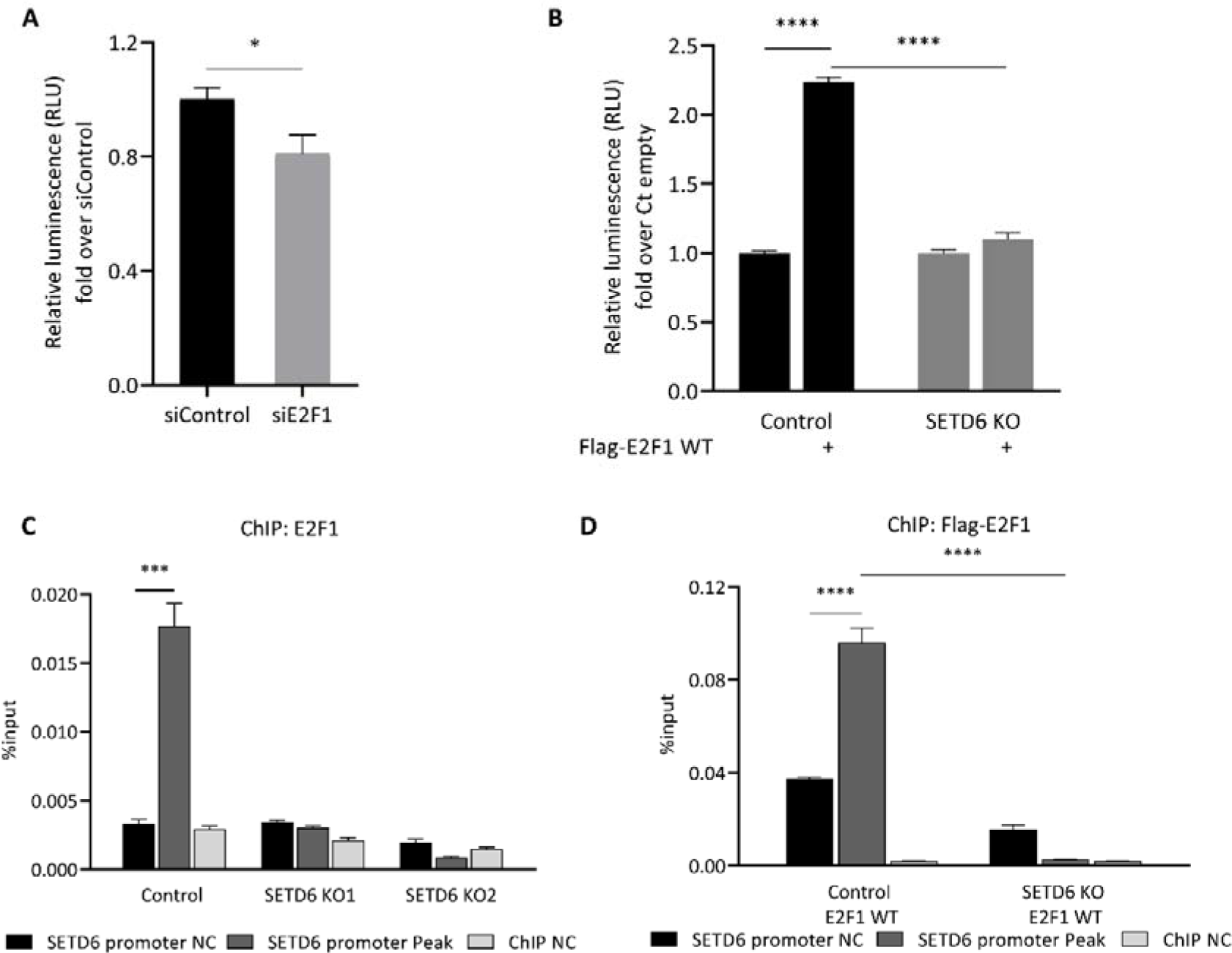
E2F1 activates SETD6 transcription in a SETD6 dependent manner. **(A)** Dual-luciferase assay in DU145 cells transfected with siRNA targeting endogenous E2F and the full promotor region of SETD6 cloned to pGL3-plasmid (positions -650 to +150 from TSS). 24 hours post-transfection, the whole cell lysates were subjected to dual-luciferase assay (Promega). Luminescence was measured by Tecan Infinite M200 plate reader. Relative luminescence was calculated after normalization of the firefly luciferase signal over Renilla luciferase control. **(B)** Same as in A with over expression of Empty or Flag-E2F1 WT in control and SETD6 KO cells. Values are fold change over siControl. Error bars are s.d. statistical analysis was performed for 3 experimental repeats. *p≤0.03, ****p≤0.0001. **(C)** Chromatin immunoprecipitation (ChIP) assay in control and SETD6 KO DU145 cells (two independent SETD6 gRNAs). **(D)** Same as in C with overexpression of Flag-E2F1 WT in control or SETD6 KO cells. 24h after transfection the chromatin fraction of the cells were immunoprecipitated with endogenous E2F1 antibody (C) or Flag antibody (D). The bound DNA was purified and amplified by qPCR using specific primers to SETD6 gene promoter regions and a distal n.c. region. Graphs show the percentage input of the quantified DNA. Error bars are s.e.m. Statistical analysis was performed for 4 experimental repeats. ***p≤0.0002, ****p≤0.0001.

To further validate this hypothesis, we performed chromatin immunoprecipitation followed by qPCR analysis (ChIP-qPCR) to compare the occupancy of the endogenous E2F1 or overexpressed Flag-E2F1 WT in DU145 control and SETD6 KO cells. ChIP-qPCR confirmed E2F1 binding to the specific region located in SETD6 promoter correlating with the peak observed in ChIP-seq data (Figure 2C and 2D). The enrichment of endogenous E2F1, as well as of the overexpressed Flag-E2F1, was significantly lower at all of the tested regions in the absence of endogenous SETD6. This data raised the hypothesis that SETD6 and E2F1 are linked and might have a functional cellular crosstalk between them in prostate cancer.

### SETD6 methylates E2F1 *in-vitro* and in cells

We first investigated the physical interaction between E2F1 and SETD6 (Supplementary Figure 3). A direct interaction between the proteins was tested in an enzyme-linked immunosorbent assay (ELISA). In these experiments, recombinant His-SUMO-E2F1, MBP-RelA as positive control, or bovine serum albumin (BSA) as negative control were immobilized on a 96-well plate, followed by incubation with recombinant glutathione S-transferase (GST) tagged SETD6 or GST. A significant direct interaction was observed between SETD6 and E2F1. Given the enzymatic activity of SETD6 and its physical interaction with E2F1 *in-vitro*, we hypothesized that SETD6 methylates E2F1.

SETD6 potential enzymatic activity on E2F1 was first tested on peptide level. To this end, 15 amino acids long peptides were synthesized on a cellulose membrane using the SPOT technology (39,40) to contain the 14 lysines within the E2F1 sequence and the corresponding K to A mutants. The RelA peptide (7), located in spot A1 served as positive control. The peptide arrays were then subjected to an *in-vitro* methylation reaction using recombinant SETD6 and ^3^H-AdoMet as the methyl group donor. Interestingly, we discovered a strong methylation signal on the E2F1 111-125 peptide containing residues K117 and K120 (spot A3). Spot A4 only containing K117 showed reduced methylation. The methylation signal was completely lost if the lysine in the peptide, corresponding to K117 in E2F1, was exchanged by alanine (spots B3 and B4), strongly implying that K117 is the primary methylation site of SETD6 in E2F1 and K120 supports the methylation (Figure 3A). To validate this finding and to examine the state of methylation, we repeated the same experiments with un-modified (WT), Kme1, Kme2 and Kme3 E2F1 (aa 111-125) peptides. On this array, we only observed a methylation signal at the WT sequence but not in the modified peptides (Figure 3B) suggesting that at the peptide level, SETD6 mono-methylates E2F1 specifically at K117. To test if K117 in full-length E2F1 is methylated by SETD6, we cloned, expressed and purified full-length WT and K117R mutant E2F1 and SETD6 (Figure 3C). The *in-vitro* methylation assay presented in Figure 3D, demonstrates that SETD6 methylates E2F1 WT but no signal was observed when E2F1 K117R mutant was used. These results indicate that SETD6 methylates E2F1 specifically at K117 *in-vitro*.

**Figure 3.**
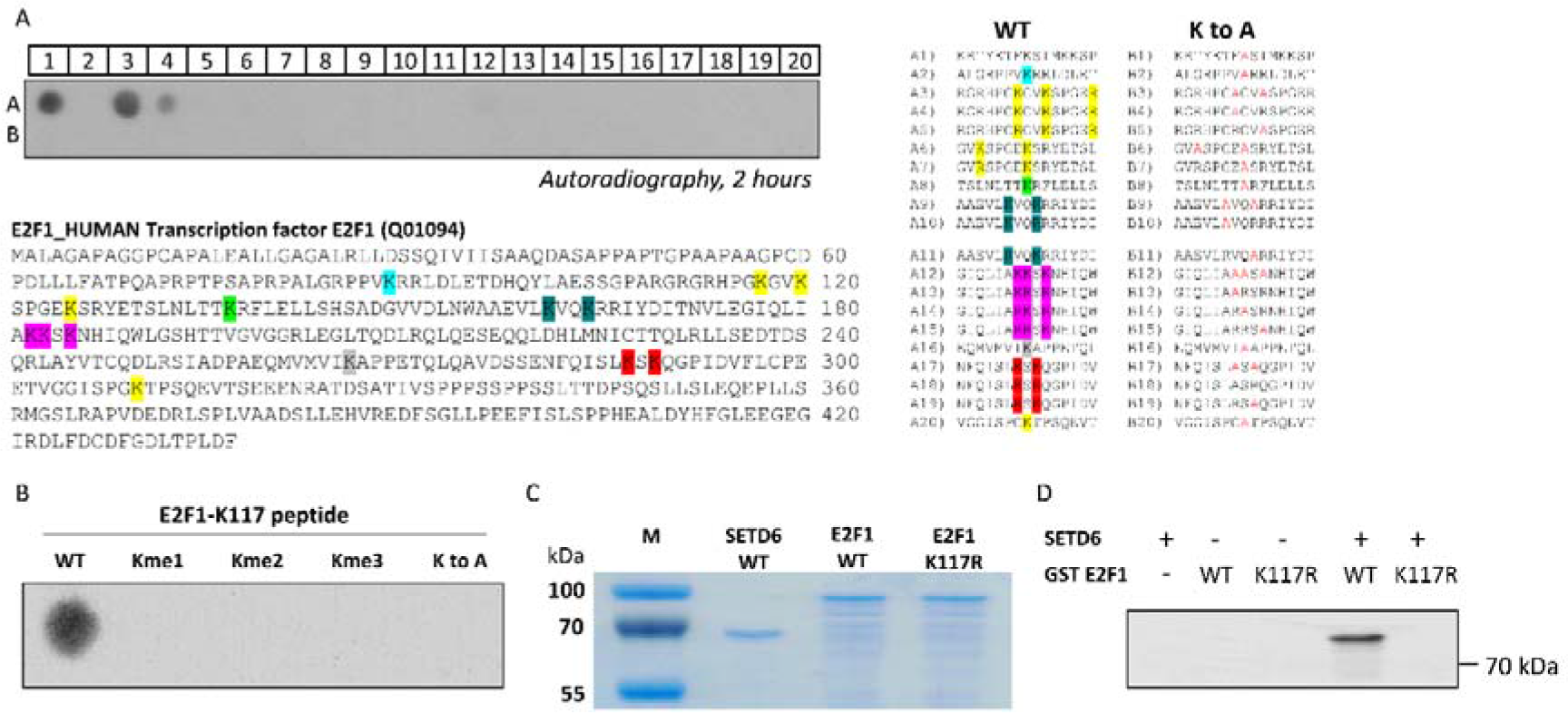
SETD6 methylates E2F1 *in-vitro* at K117 at peptide and protein level. **(A)** Methylation of a peptide SPOT array for determination of the target lysine of SETD6 on E2F1. 15 aa long peptides containing all lysine residues of E2F1 and variants in which individual lysine residues were replaced by alanine were synthesized on a peptide SPOT array, and incubated with recombinant SETD6 in the presence of radioactively labelled AdoMet. The RelA peptide (in spot A1) served as positive control, while the mutated RelA K310A in spot B1 served as the negative control. Peptide sequences of the individual spots are presented on the right. **(B)** Same experiment as described in panel A was performed with a peptide array containing unmodified (WT), or modified (Kme1, Kme2, Kme3 and a K to A mutated) E2F1 K117 peptides (aa 111-125, same as in spot A3). **(C)** Coomassie stain of purified His-SETD6, GST-E2F1 WT and GST-E2F1 K117R mutant to show equal loading of substrate proteins and enzyme used for *in-vitro* methylation experiments. **(D)** *In-vitro* methylation assay of GST-E2F1 (WT and the mutant K117R) by recombinant His-SETD6 in the presence of radioactively labelled AdoMet. The transferred radioactively labelled methyl groups were detected by autoradiography after different exposure times.

To validate that E2F1 is methylated at K117 in cells, we immunoprecipitated overexpressed Flag-E2F1 WT and the K117R-mutant from DU145 using a pan-Kme1antibody (Figure 4A). While a strong methylation was observed for E2F1 WT, the methylation of E2F1 K117R in DU145 decreased. We next purchased a custom made site-specific antibody for E2F1 K117me1. Validation of the antibody specificity in a dot-blot experiment shows that it specifically recognizes mono-methylated E2F1 K117, but not the un-modified and the Kme2 and Kme3 peptides (Figure 4B). We next utilized this antibody for an *in-vitro* methylation assay with recombinant proteins showing that E2F1 WT but not the K117R mutant is methylated in the presence of recombinant SETD6 (Figure 4C). Using this antibody, we confirmed that the methylation of over-expressed E2F1 depends on the enzymatic activity of SETD6 as no methylation was observed when the catalytic Y285A of SETD6 (7) was expressed (Figure 4D). In a complementary experiment, we found that SETD6 methylates E2F1 in control cell but not when E2F1 K117R mutant was used or in SETD6 KO cells (Figure 4E). Taken together, these results indicate that E2F1 is methylated by SETD6 *in-vitro* and in cells and lysine 117 is the primary methylation site.

**Figure 4.**
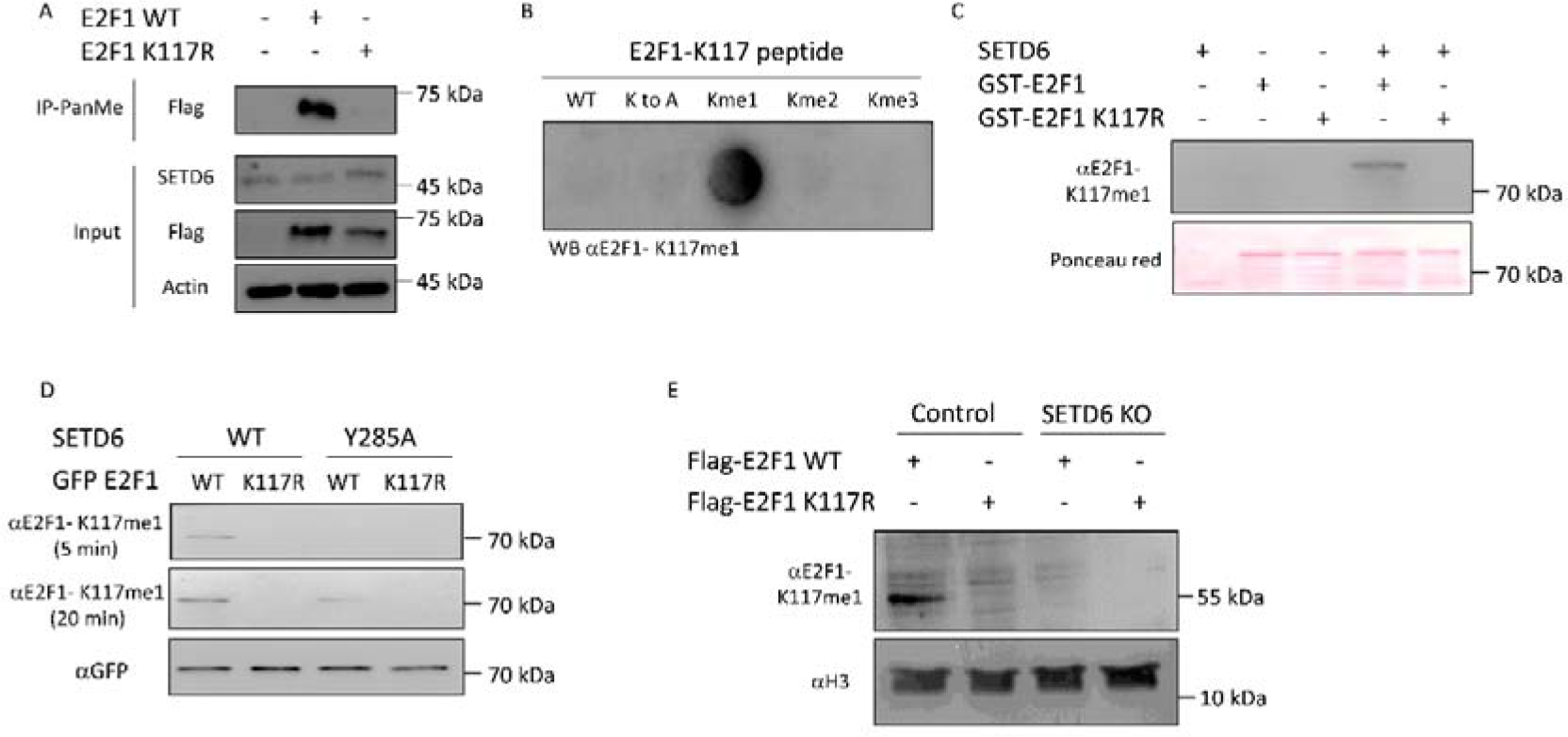
SETD6 methylates E2F1 at K117 in cells. **(A)** DU145 cells were transfected with Flag-E2F1 WT or the respective K117R-mutant. After 24 hours, whole cell lysates were immunoprecipitated with Pan-Kme1 antibody, followed by WB analysis with indicated antibodies. **(B)** For validation of the specificity of the custom made E2F1-K117me1 antibody, a peptide SPOT array with 15 aa long unmodified (WT), or modified (Kme1, Kme2, Kme3 and a K to A mutated) E2F1 peptides (aa 111-125) was incubated with the antibody and binding analyzed by a secondary antibody and ECL. **(C)** *In-vitro* methylation assay GST-E2F1 (WT and the mutant K117R) by recombinant His-SETD6 using unlabelled AdoMet as cofactor. After methylation WB analysis was performed using the E2F1-K117me1 antibody. **(D)** DU145 cells were transfected with the indicated plasmids. 72 hours post-transfection, the methylation levels of EGFP-fused E2F1 were assessed by WB using the E2F1-K117me1 antibody. WB against GFP was included as input control. **(E)** DU145 control and SETD6 KO cells were transfected with the indicated plasmids. After 24 hours the whole protein lysate was analyzed by WB with the E2F1-K117me1 antibody. WB against H3 was included as input loading control.

### E2F1 methylation at K117 regulates SETD6 promoter activation

Having demonstrated that E2F1 binds the SETD6 promoter and activates its transcription, we hypothesized that there could be a molecular feedback mechanism by which the methylation of E2F1 at K117 by SETD6 may affect the transcription of SETD6. To address this hypothesis, we first tested the activity of the recombinant SETD6 promotor driving luciferase expression in cells over-expressing E2F1 WT or E2F1 K117R mutant. A significant increase in the luciferase activity was observed in cells overexpressing WT compared to the K117R-mutant (Figure 5A). Consistent with these results, a significant elevation in endogenous SETD6 mRNA expression level was observed in cells overexpressing WT E2F1 compared to the E2F1 K117R mutant (Figure 5B). To validate these results, we performed ChIP-qPCR experiments to test the SETD6 promoter occupancy of E2F1 in DU145 control and SETD6 KO cells overexpressing Flag-E2F1 WT or Flag-E2F1 K117R mutant, using the specific E2F1 K117me1 antibody for pulldown (Figure 5C). The methylated E2F1 was identified more abundantly at the predicted region of SETD6 promoter in overexpressed Flag-E2F1 WT DU145 control cells than in Flag-E2F1 K117R or SETD6 KO cells (Figure 5C). These data provide evidence that E2F1 is present at the promoter of SETD6 and specifically, methylated E2F1 at K117 is more inclined to bind to this site. Collectively our results suggest that methylated E2F1 binds the SETD6 promoter and activates its expression in a SETD6 and a methylation dependent manner (Figure 5D).

**Figure 5.**
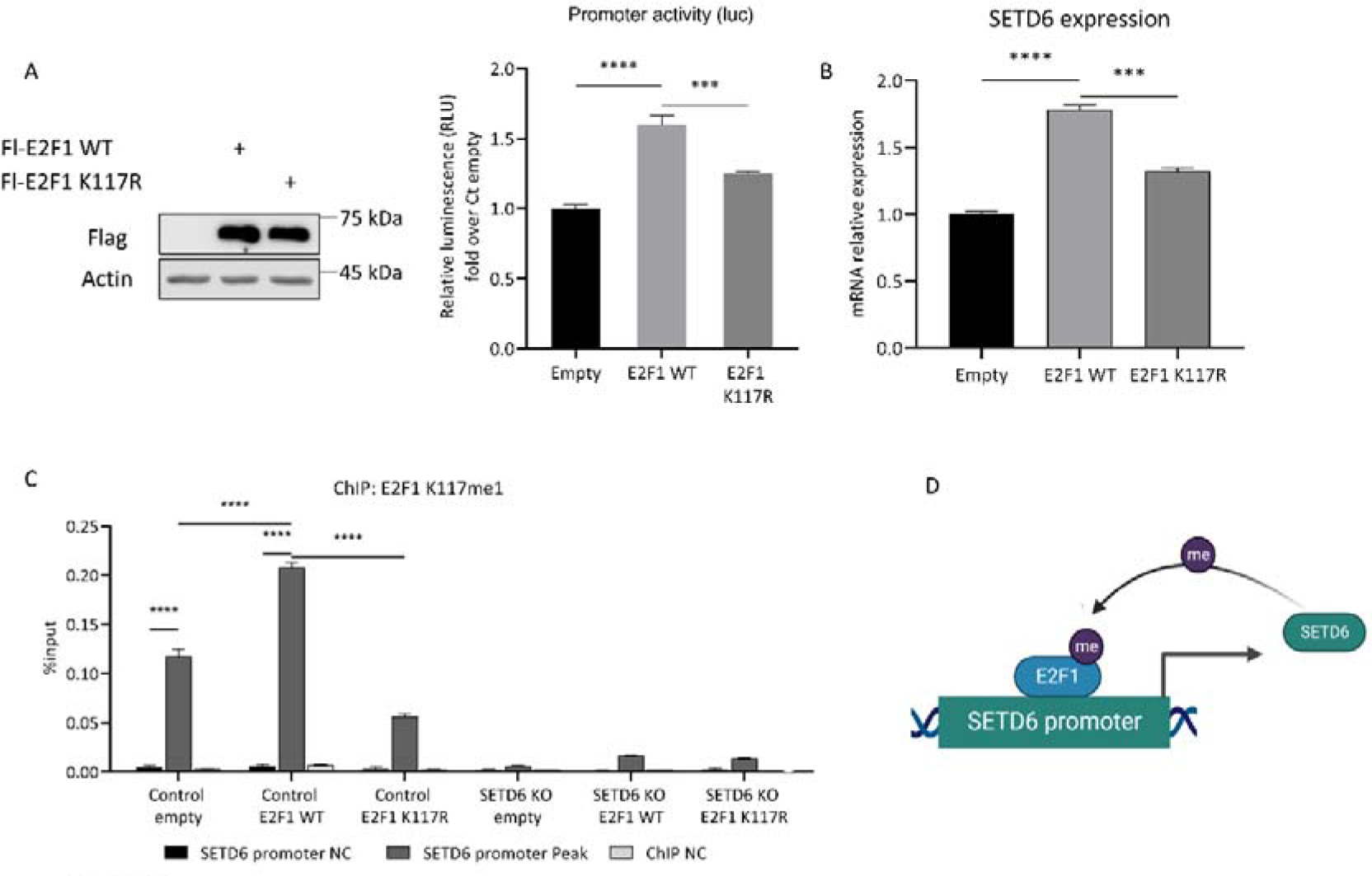
E2F1 regulates SETD6 mRNA levels in a SETD6 dependent manner. **(A)** Dual-luciferase assay 24h after transfection with empty plasmid, Flag-E2F1 WT or Flag-E2F1 K117R mutant and the full-length SETD6 promoter luciferase construct. Values are fold change over empty control. E2F1 protein levels were assessed by WB (left). **(B)** DU145 cells were transfected with empty plasmid, Flag-E2F1 WT or Flag-E2F1 K117R mutant. SETD6 mRNA expression levels were measured using RT-qPCR. mRNA expression levels were normalized to mRNA expression levels of GAPDH housekeeping gene. **(C)** Chromatin immunoprecipitation (ChIP) assay 24h after transfection with empty plasmid, Flag-E2F1 WT or Flag-E2F1 K117R mutant. The chromatin fractions of the cells were immunoprecipitated with anti-E2F1 K117me1. Error bars are s.e.m. Statistical analysis was performed for 5 experimental repeats. **p≤0.002, ***p≤0.0002, ****p≤0.00001. **(D)** Graphical model of our findings. E2F1 methylation by SETD6 regulates the mRNA expression of SETD6 in a positive feedback mechanism.

## Discussion

In recent years, lysine methylation has been identified as an integral part of cellular biology, and a key regulator of physiological and pathological processes in the cell. However, only a fraction of methylation events in the human proteome has been characterized. Here we provide further insights into the lysine methylation field by the identification of E2F1 as a new SETD6 substrate which in turn regulates the expression of SETD6 in a methylation dependent manner. This SETD6-mediated methylation of E2F1 may have implications for the progression of diseases, including cancer. We may envision that the positive feedback mechanism which is mediated by E2F1 methylation is not only important for the tight regulation of SETD6 transcription but also for SETD6 enzymatic activity to methylate other downstream substrates. Such a mechanism will ensure maintenance of the steady state level of SETD6 to allow proper regulation of cellular pathways in which SETD6 is involved (7-12,15).

E2F1 activity has been shown to be regulated through extensive post-translational modifications (41–44). Interestingly, E2F1 is also subjected to acetylation (45,46), arginine methylation (47) and NEDDylation(48) at K117, the exact same residue we have identified in this study to be methylated by SETD6. These modifications are located near the DNA binding domain of E2F1. E2F1 is acetylated at K117, K120 and K125 in response to DNA damage (49). This modification was shown to increase E2F1 stability, DNA binding affinity (45), and create a binding motif for the bromodomains of the p300/KAT3B and CBP/KAT3A acetyltransferases. Lysine residues 117, 120, 125, 182, 183 and 185 are required for efficient NEDDylation of E2F1 by ubiquitin-like modifier NEDD8. Experimental evidence suggests that K185 is particularly important for this PTM (48). NEDDylation results in decreased E2F1 stability, lower transcriptional activity, and slower cell growth. This specific post-translational modification is regulated by SETD7-mediated methylation of E2F1 at K185, in response to DNA damage (50). This modification attenuates the level of E2F1 expression, by inhibition of acetylation and phosphorylation of the protein at distant residues and, simultaneously, by stimulation of poly-ubiquitination and subsequent degradation of E2F1 by 26S proteasome (41,50). Yet, E2F1 forms a negative regulatory loop with SETD7, as it was demonstrated that SETD7 co-activates E2F1-dependent transcription of CCNE1 gene, thus promoting cell proliferation, through successful exit from the G1/S checkpoint arrest (51). While it can be assumed that methylation of K117 prevents other types of modifications at this residue, future studies are required to deeply assess if there is a physiological and pathological link between these different modifications of E2F1 and the methylation of K117 by SETD6.

We have previously shown that BRD4 methylation by SETD6 at K99 regulates the recruitment of E2F1 to chromatin to selectively regulate the expression genes involved in protein translation. Once BRD4 is methylated, the recruitment of E2F1 to translation-related target genes is inhibited in a SETD6 and K99 methylation dependent manner (10). Now, with the discovery that E2F1 is also methylated by SETD6, an intriguing working hypothesis that should be examined in the future is to test if E2F1 has to be methylated at K117 in order to control its association with the DNA and its genomic distribution.

Our results indicate that the methylation of K117 on E2F1 by SETD6 has an impact on E2F1’s binding to the promoter region of the SETD6 gene, thereby modulating SETD6 gene expression. Specifically, methylation of E2F1 at K117 results in an increase in E2F1’s enrichment at the SETD6 promoter and subsequently augments transcription of the SETD6 promoter, leading to elevated levels of SETD6 mRNA expression. We provide evidence for the existence of SETD6-E2F1 feedback loop. The characterization of feedbacks mechanisms have an enormous contribution for our understanding of cellular signaling pathways and network dynamics in many physiological and pathological scenarios (52). This include also feedback mechanisms which are mediated by E2F1 such in the NFKB (53) and the KRAS (54,55) signaling pathways. Therefore, it is not unusual that methylation by SETD6 directs E2F1 transcriptional activity to regulate SETD6 mRNA transcription and may have a regulatory role to govern functional dynamics of cellular processes.

The transcription factor E2F1 has many roles in the regulation of diverse oncogenic-related cellular pathways and phenotypes in several types of cancers (56–59). In prostate cancer specifically, E2F1 was shown to act in a dichotomic manner in several oncogenic processes (60). However, the regulation of E2F1 transcriptional activity is still poorly understood. In this paper, using biochemical and molecular biology approaches, we discovered how methylation of E2F1 by SETD6, creates a feedback loop in the regulation of SETD6 mRNA expression levels.

Further research is required to determine how SETD6 mRNA levels affect the malignant progression of prostate cancer. Through the identification of the precise mechanisms by which SETD6-mediated methylation of E2F1 modulates gene expression, it will become possible to develop novel approaches for the precise and targeted modulation of gene activity. This holds significant promise for the development of new therapies for a diverse range of diseases, including cancer and other disorders characterized by abnormal gene expression patterns.

## Funding

This work was supported by a joint grant of the DFG to AJ and DL (JE 252/38-1). DL was also supported by The Israel Science Foundation (285/14 and 262/18), The Israeli Cancer Research Foundation Israel (ICRF), and from the Israel Cancer Association.

## Acknowledgments

We thank the Levy and Jeltsch lab for technical assistance and helpful discussions.

## Author Contribution

DL, MK, SW and AJ conceived the work and were involved experimental design. MK, GU and SW performed most of the experiments. DL, MK, GU, SW and AJ were involved in data analysis and interpretation. NL and MF assisted in cloning, protein expression and purification and performed the ELISA experiment. MK, DL and AJ wrote the paper. All authors read and approved the final manuscript.

## Conflict of Interest

The authors declare that they have no conflict of interest.

---

**Supplementary Figure 1.**
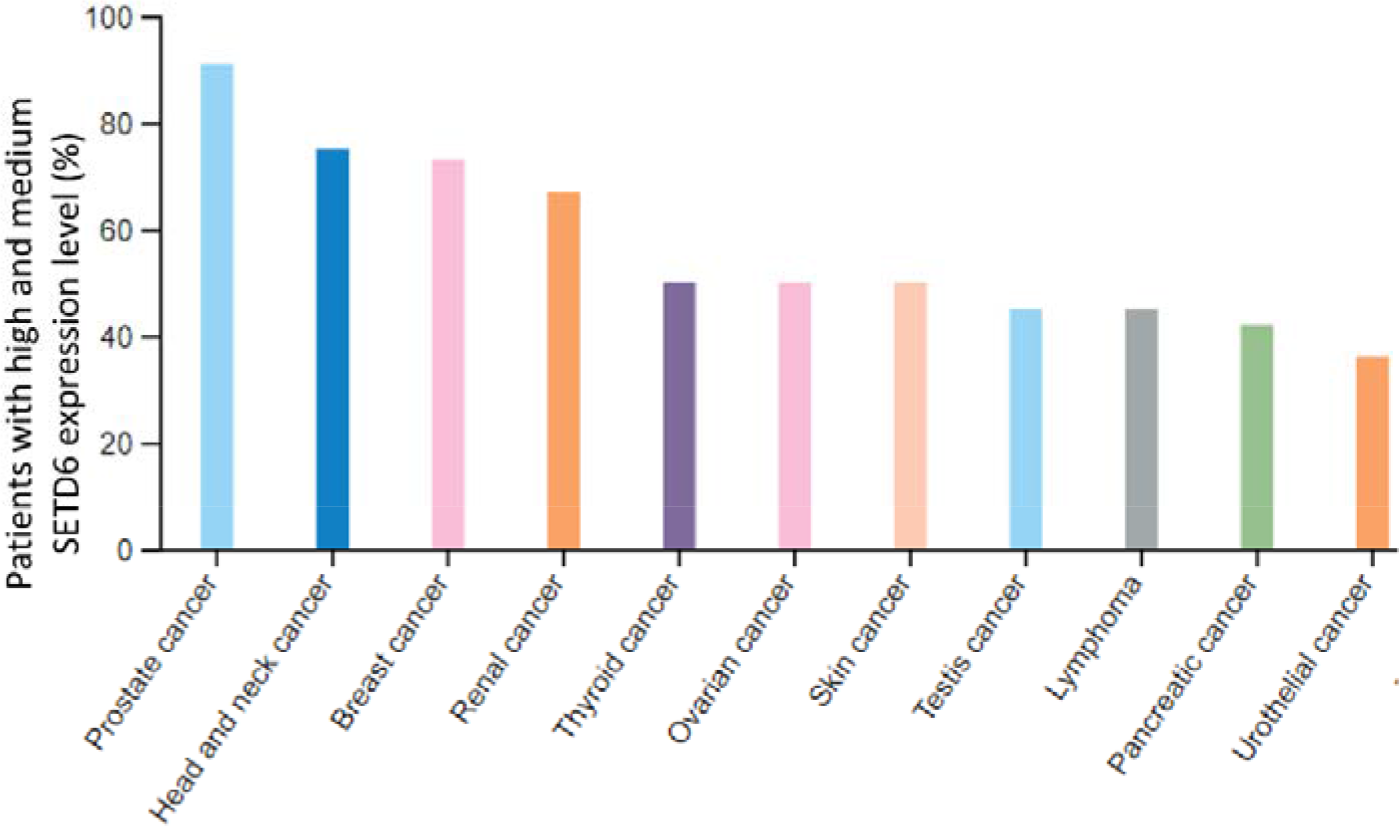
Expression levels of SETD6 in various cancer. The graphs were generated using the Human Protein Atlas resource (https://www.proteinatlas.org/). For each cancer, color-coded bars indicate the percentage of patients (maximum 12 patients) with high and medium SETD6 protein expression level.

**Supplementary Figure 2.**
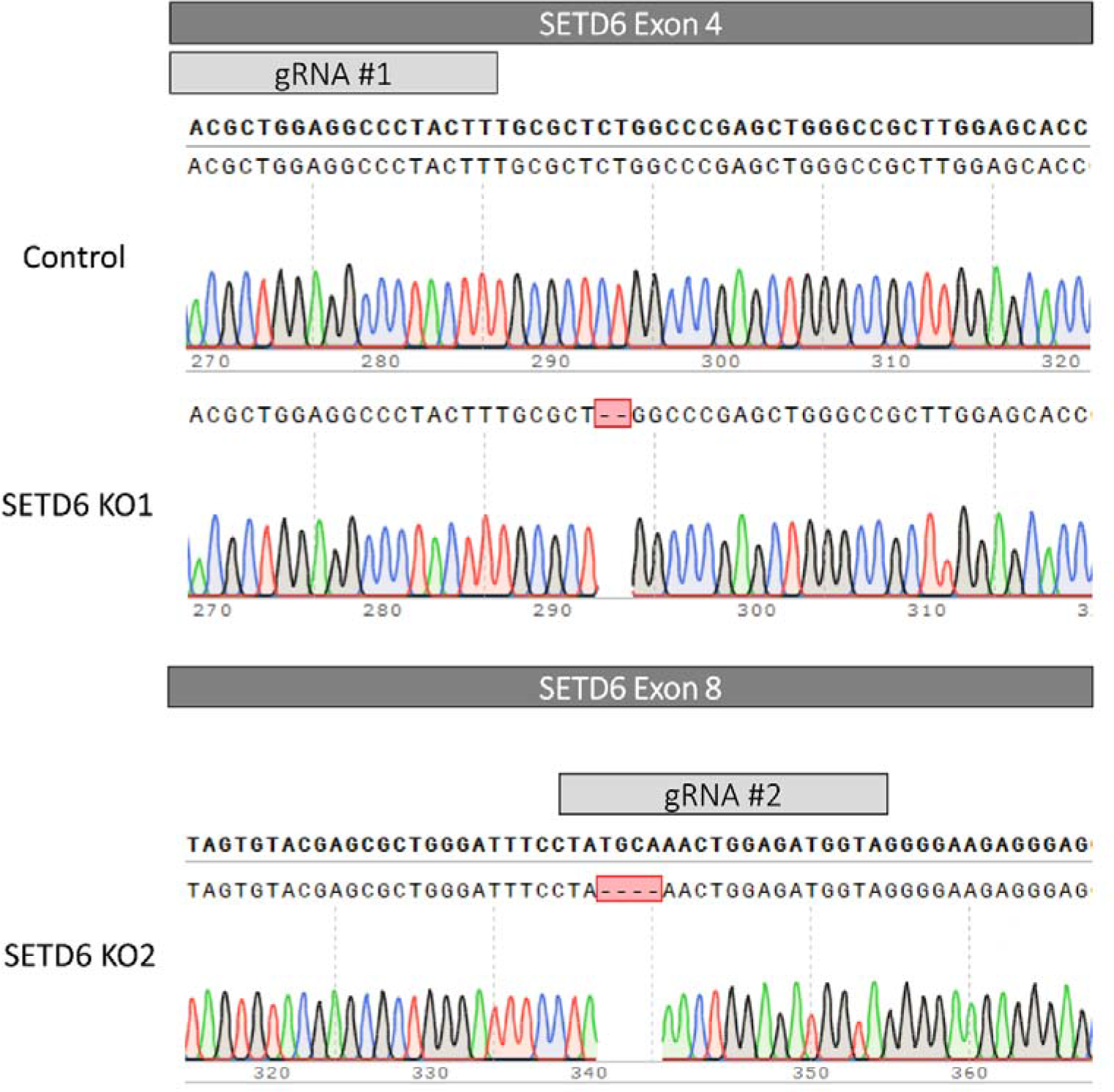
SETD6 KO validation. DU145 control and SETD6 knock-out (SETD6 KO) were generated using the CRISPR/Cas9 system, following single-clone selection. Chromatograms of Sanger sequencing of control cells and two SETD6 KO cells generated from two independent gRNAs targeted for SETD6 exon 4 and exon 8.

**Supplementary Figure 3.**
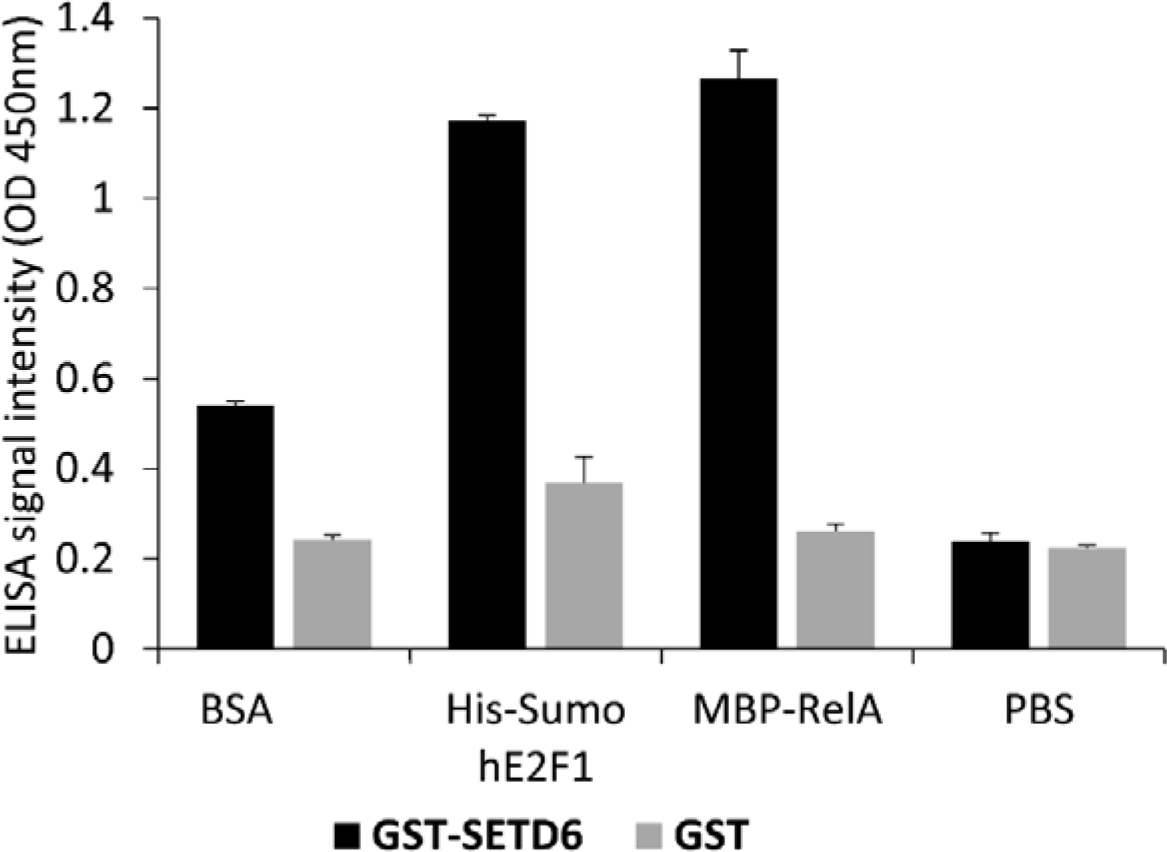
ELISA assay showing the interaction assessment between recombinant SETD6 and E2F1. His-SUMO-E2F1, MBP-RelA or BSA were bound to the surface of a 96-well plate, incubated with GST-SETD6, or GST protein as negative control, and binding was analyzed with anti-GST antibody. RelA served as positive control for interaction with SETD6, while BSA served as negative control. PBS served as background control. Error bars are s.d. based on 3 replicas.

**Supplementary Table 1.**
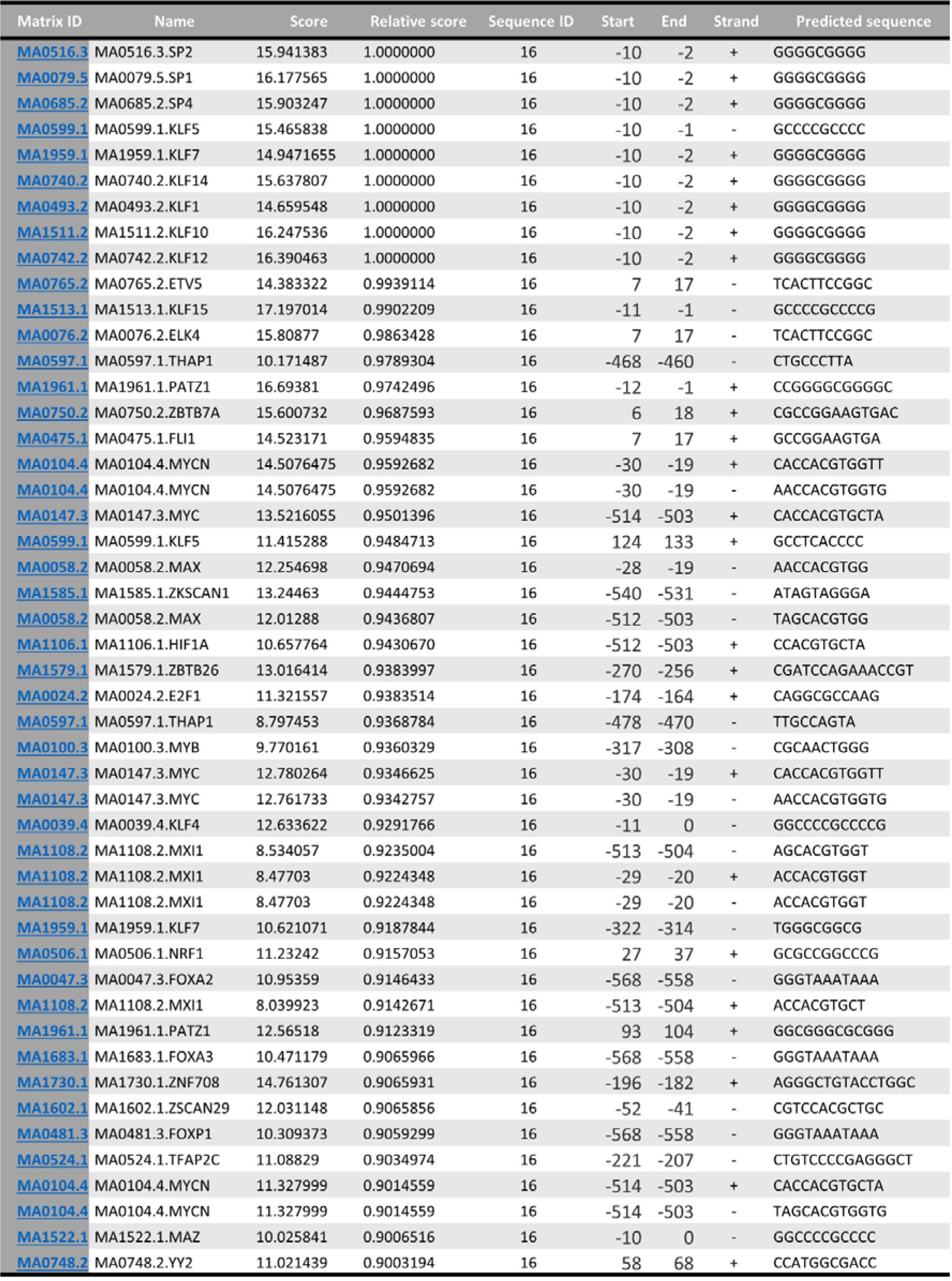
E2F1 predicted binding sites. E2F1 binding sites at the SETD6 promoter sequence, as predicted by the JASPAR database (https://jaspar.genereg.net/) with relative profile score threshold of 90%. The prediction is based on curated transcription factors binding profiles as position frequency matrices (PFMs).

## Notes

### Competing Interest Statement

The authors have declared no competing interest.

### Summary of Updates

authoer order was changed

## References

1. Alam, H., Gu, B., and Lee, M. G. (2015) Histone methylation modifiers in cellular signaling pathways. Cell Mol Life Sci 72, 4577–4592

2. Kudithipudi, S., and Jeltsch, A. (2016) Approaches and Guidelines for the Identification of Novel Substrates of Protein Lysine Methyltransferases. Cell Chem Biol 23, 1049–1055

3. Morera, L., Lubbert, M., and Jung, M. (2016) Targeting histone methyltransferases and demethylases in clinical trials for cancer therapy. Clin Epigenetics 8, 57

4. Del Rizzo, P. A., and Trievel, R. C. (2011) Substrate and product specificities of SET domain methyltransferases. Epigenetics 6, 1059–1067

5. Weil, L. E., Shmidov, Y., Kublanovsky, M., Morgenstern, D., Feldman, M., Bitton, R., and Levy, D. (2018) Oligomerization and Auto-methylation of the Human Lysine Methyltransferase SETD6. J Mol Biol 430, 4359–4368

6. O’Neill, D. J., Williamson, S. C., Alkharaif, D., Monteiro, I. C., Goudreault, M., Gaughan, L., Robson, C. N., Gingras, A. C., and Binda, O. (2014) SETD6 controls the expression of estrogen-responsive genes and proliferation of breast carcinoma cells. Epigenetics 9, 942–950

7. Levy, D., Kuo, A. J., Chang, Y., Schaefer, U., Kitson, C., Cheung, P., Espejo, A., Zee, B. M., Liu, C. L., Tangsombatvisit, S., Tennen, R. I., Kuo, A. Y., Tanjing, S., Cheung, R., Chua, K. F., Utz, P. J., Shi, X., Prinjha, R. K., Lee, K., Garcia, B. A., Bedford, M. T., Tarakhovsky, A., Cheng, X., and Gozani, O. (2011) Lysine methylation of the NF-kappaB subunit RelA by SETD6 couples activity of the histone methyltransferase GLP at chromatin to tonic repression of NF-kappaB signaling. Nat Immunol 12, 29–36

8. Admoni-Elisha, L., Elbaz, T., Chopra, A., Shapira, G., Bedford, M. T., Fry, C. J., Shomron, N., Biggar, K., Feldman, M., and Levy, D. (2022) TWIST1 methylation by SETD6 selectively antagonizes LINC-PINT expression in glioma. Nucleic Acids Res 50, 6903–6918

9. Admoni-Elisha, L., Abaev-Schneiderman, E., Cohn, O., Shapira, G., Shomron, N., Feldman, M., and Levy, D. (2022) Structure-function conservation between the methyltransferases SETD3 and SETD6. Biochimie 200, 27–35

10. Vershinin, Z., Feldman, M., Werner, T., Weil, L. E., Kublanovsky, M., Abaev-Schneiderman, E., Sklarz, M., Lam, E. Y. N., Alasad, K., Picaud, S., Rotblat, B., McAdam, R. A., Chalifa-Caspi, V., Bantscheff, M., Chapman, T., Lewis, H. D., Filippakopoulos, P., Dawson, M. A., Grandi, P., Prinjha, R. K., and Levy, D. (2021) BRD4 methylation by the methyltransferase SETD6 regulates selective transcription to control mRNA translation. Sci Adv 7

11. Vershinin, Z., Feldman, M., and Levy, D. (2020) PAK4 methylation by the methyltransferase SETD6 attenuates cell adhesion. Sci Rep 10, 17068

12. Feldman, M., Vershinin, Z., Goliand, I., Elia, N., and Levy, D. (2019) The methyltransferase SETD6 regulates Mitotic progression through PLK1 methylation. Proc Natl Acad Sci U S A 116, 1235–1240

13. Kublanovsky, M., Aharoni, A., and Levy, D. (2018) Enhanced PKMT-substrate recognition through non active-site interactions. Biochem Biophys Res Commun 501, 1029–1033

14. Feldman, M., and Levy, D. (2018) Peptide inhibition of the SETD6 methyltransferase catalytic activity. Oncotarget 9, 4875–4885

15. Martin-Morales, L., Feldman, M., Vershinin, Z., Garre, P., Caldes, T., and Levy, D. (2017) SETD6 dominant negative mutation in familial colorectal cancer type X. Hum Mol Genet 26, 4481–4493

16. Cohn, O., Chen, A., Feldman, M., and Levy, D. (2016) Proteomic analysis of SETD6 interacting proteins. Data in brief 6, 799–802

17. Vershinin, Z., Feldman, M., Chen, A., and Levy, D. (2016) PAK4 Methylation by SETD6 Promotes the Activation of the Wnt/beta-Catenin Pathway. J Biol Chem 291, 6786–6795

18. Chen, A., Feldman, M., Vershinin, Z., and Levy, D. (2016) SETD6 is a negative regulator of oxidative stress response. Biochim Biophys Acta 1859, 420–427

19. Chang, Y., Levy, D., Horton, J. R., Peng, J., Zhang, X., Gozani, O., and Cheng, X. (2011) Structural basis of SETD6-mediated regulation of the NF-kB network via methyl-lysine signaling. Nucleic Acids Res 39, 6380–6389

20. Webb, W. M., Irwin, A. B., Pepin, M. E., Henderson, B. W., Huang, V., Butler, A. A., Herskowitz, J. H., Wende, A. R., Cash, A. E., and Lubin, F. D. (2020) The SETD6 Methyltransferase Plays an Essential Role in Hippocampus-Dependent Memory Formation. Biol Psychiatry 87, 577–587

21. Jose, L., Androphy, E. J., and DeSmet, M. (2022) SETD6 Regulates E2-Dependent Human Papillomavirus Transcription. J Virol 96, e0129522

22. Xu, J., Zhou, H., Luo, Z., Chen, J., and Liu, M. (2023) Investigating the functional role of SETD6 in lung adenocarcinoma. BMC Cancer 23, 18

23. Mukherjee, N., Cardenas, E., Bedolla, R., and Ghosh, R. (2017) SETD6 regulates NF-kappaB signaling in urothelial cell survival: Implications for bladder cancer. Oncotarget 8, 15114–15125

24. Dimova, D. K., and Dyson, N. J. (2005) The E2F transcriptional network: old acquaintances with new faces. Oncogene 24, 2810–2826

25. Bracken, A. P., Ciro, M., Cocito, A., and Helin, K. (2004) E2F target genes: unraveling the biology. Trends Biochem Sci 29, 409–417

26. Polager, S., Kalma, Y., Berkovich, E., and Ginsberg, D. (2002) E2Fs up-regulate expression of genes involved in DNA replication, DNA repair and mitosis. Oncogene 21, 437–446

27. Zhu, W., Giangrande, P. H., and Nevins, J. R. (2004) E2Fs link the control of G1/S and G2/M transcription. EMBO J 23, 4615–4626

28. Giangrande, P. H., Zhu, W., Rempel, R. E., Laakso, N., and Nevins, J. R. (2004) Combinatorial gene control involving E2F and E Box family members. EMBO J 23, 1336–1347

29. Araki, K., Nakajima, Y., Eto, K., and Ikeda, M. A. (2003) Distinct recruitment of E2F family members to specific E2F-binding sites mediates activation and repression of the E2F1 promoter. Oncogene 22, 7632–7641

30. Le Cam, L., Polanowska, J., Fabbrizio, E., Olivier, M., Philips, A., Ng Eaton, E., Classon, M., Geng, Y., and Sardet, C. (1999) Timing of cyclin E gene expression depends on the regulated association of a bipartite repressor element with a novel E2F complex. EMBO J 18, 1878–1890

31. Giangrande, P. H., Hallstrom, T. C., Tunyaplin, C., Calame, K., and Nevins, J. R. (2003) Identification of E-box factor TFE3 as a functional partner for the E2F3 transcription factor. Mol Cell Biol 23, 3707–3720

32. Chung, C. W., Coste, H., White, J. H., Mirguet, O., Wilde, J., Gosmini, R. L., Delves, C., Magny, S. M., Woodward, R., Hughes, S. A., Boursier, E. V., Flynn, H., Bouillot, A. M., Bamborough, P., Brusq, J. M., Gellibert, F. J., Jones, E. J., Riou, A. M., Homes, P., Martin, S. L., Uings, I. J., Toum, J., Clement, C. A., Boullay, A. B., Grimley, R. L., Blandel, F. M., Prinjha, R. K., Lee, K., Kirilovsky, J., and Nicodeme, E. (2011) Discovery and characterization of small molecule inhibitors of the BET family bromodomains. J Med Chem 54, 3827–3838

33. Ramos-Montoya, A., Lamb, A. D., Russell, R., Carroll, T., Jurmeister, S., Galeano-Dalmau, N., Massie, C. E., Boren, J., Bon, H., Theodorou, V., Vias, M., Shaw, G. L., Sharma, N. L., Ross-Adams, H., Scott, H. E., Vowler, S. L., Howat, W. J., Warren, A. Y., Wooster, R. F., Mills, I. G., and Neal, D. E. (2014) HES6 drives a critical AR transcriptional programme to induce castration-resistant prostate cancer through activation of an E2F1-mediated cell cycle network. EMBO Mol Med 6, 651–661

34. Barfeld, S. J., Urbanucci, A., Itkonen, H. M., Fazli, L., Hicks, J. L., Thiede, B., Rennie, P. S., Yegnasubramanian, S., DeMarzo, A. M., and Mills, I. G. (2017) c-Myc Antagonises the Transcriptional Activity of the Androgen Receptor in Prostate Cancer Affecting Key Gene Networks. EBioMedicine 18, 83–93

35. Bert, S. A., Robinson, M. D., Strbenac, D., Statham, A. L., Song, J. Z., Hulf, T., Sutherland, R. L., Coolen, M. W., Stirzaker, C., and Clark, S. J. (2013) Regional activation of the cancer genome by long-range epigenetic remodeling. Cancer Cell 23, 9–22

36. Liu, Y., Yu, S., Dhiman, V. K., Brunetti, T., Eckart, H., and White, K. P. (2017) Functional assessment of human enhancer activities using whole-genome STARR-sequencing. Genome Biol 18, 219

37. Robinson, J. T., Thorvaldsdottir, H., Turner, D., and Mesirov, J. P. (2023) igv.js: an embeddable JavaScript implementation of the Integrative Genomics Viewer (IGV). Bioinformatics 39

38. Castro-Mondragon, J. A., Riudavets-Puig, R., Rauluseviciute, I., Lemma, R. B., Turchi, L., Blanc-Mathieu, R., Lucas, J., Boddie, P., Khan, A., Manosalva Perez, N., Fornes, O., Leung, T. Y., Aguirre, A., Hammal, F., Schmelter, D., Baranasic, D., Ballester, B., Sandelin, A., Lenhard, B., Vandepoele, K., Wasserman, W. W., Parcy, F., and Mathelier, A. (2022) JASPAR 2022: the 9th release of the open-access database of transcription factor binding profiles. Nucleic Acids Res 50, D165–D173

39. Weirich, S., and Jeltsch, A. (2022) Specificity Analysis of Protein Methyltransferases and Discovery of Novel Substrates Using SPOT Peptide Arrays. Methods Mol Biol 2529, 313–325

40. Frank, R. (2002) The SPOT-synthesis technique. Synthetic peptide arrays on membrane supports--principles and applications. J Immunol Methods 267, 13–26

41. Moiseeva, T. N., Bottrill, A., Melino, G., and Barlev, N. A. (2013) DNA damage-induced ubiquitylation of proteasome controls its proteolytic activity. Oncotarget 4, 1338–1348

42. Real, S., Espada, L., Espinet, C., Santidrian, A. F., and Tauler, A. (2010) Study of the in vivo phosphorylation of E2F1 on Ser403. Biochim Biophys Acta 1803, 912–918

43. Galbiati, L., Mendoza-Maldonado, R., Gutierrez, M. I., and Giacca, M. (2005) Regulation of E2F-1 after DNA damage by p300-mediated acetylation and ubiquitination. Cell Cycle 4, 930–939

44. Berkovich, E., and Ginsberg, D. (2003) ATM is a target for positive regulation by E2F-1. Oncogene 22, 161–167

45. Manickavinayaham, S., Velez-Cruz, R., Biswas, A. K., Bedford, E., Klein, B. J., Kutateladze, T. G., Liu, B., Bedford, M. T., and Johnson, D. G. (2019) E2F1 acetylation directs p300/CBP-mediated histone acetylation at DNA double-strand breaks to facilitate repair. Nat Commun 10, 4951

46. Ianari, A., Gallo, R., Palma, M., Alesse, E., and Gulino, A. (2004) Specific role for p300/CREB-binding protein-associated factor activity in E2F1 stabilization in response to DNA damage. J Biol Chem 279, 30830–30835

47. Cho, E. C., Zheng, S., Munro, S., Liu, G., Carr, S. M., Moehlenbrink, J., Lu, Y. C., Stimson, L., Khan, O., Konietzny, R., McGouran, J., Coutts, A. S., Kessler, B., Kerr, D. J., and Thangue, N. B. (2012) Arginine methylation controls growth regulation by E2F-1. EMBO J 31, 1785–1797

48. Aoki, I., Higuchi, M., and Gotoh, Y. (2013) NEDDylation controls the target specificity of E2F1 and apoptosis induction. Oncogene 32, 3954–3964

49. Lin, W. C., Lin, F. T., and Nevins, J. R. (2001) Selective induction of E2F1 in response to DNA damage, mediated by ATM-dependent phosphorylation. Genes Dev 15, 1833–1844

50. Kontaki, H., and Talianidis, I. (2010) Lysine methylation regulates E2F1-induced cell death. Mol Cell 39, 152–160

51. Lezina, L., Aksenova, V., Ivanova, T., Purmessur, N., Antonov, A. V., Tentler, D., Fedorova, O., Garabadgiu, A. V., Talianidis, I., Melino, G., and Barlev, N. A. (2014) KMTase Set7/9 is a critical regulator of E2F1 activity upon genotoxic stress. Cell Death Differ 21, 1889–1899

52. Gu, W., Shen, H., Xie, L., Zhang, X., and Yang, J. (2022) The Role of Feedback Loops in Targeted Therapy for Pancreatic Cancer. Front Oncol 12, 800140

53. Ren, X., Chen, C., Luo, Y., Liu, M., Li, Y., Zheng, S., Ye, H., Fu, Z., Li, M., Li, Z., and Chen, R. (2020) lncRNA-PLACT1 sustains activation of NF-kappaB pathway through a positive feedback loop with IkappaBalpha/E2F1 axis in pancreatic cancer. Mol Cancer 19, 35

54. Chu, P. C., Yang, M. C., Kulp, S. K., Salunke, S. B., Himmel, L. E., Fang, C. S., Jadhav, A. M., Shan, Y. S., Lee, C. T., Lai, M. D., Shirley, L. A., Bekaii-Saab, T., and Chen, C. S. (2016) Regulation of oncogenic KRAS signaling via a novel KRAS-integrin-linked kinase-hnRNPA1 regulatory loop in human pancreatic cancer cells. Oncogene 35, 3897–3908

55. Chu, P. C., Kulp, S. K., Bekaii-Saab, T., and Chen, C. S. (2018) Targeting integrin-linked kinase to suppress oncogenic KRAS signaling in pancreatic cancer. Small GTPases 9, 452–456

56. Tang, Y., Jiang, L., Zhao, X., Hu, D., Zhao, G., Luo, S., Du, X., and Tang, W. (2021) FOXO1 inhibits prostate cancer cell proliferation via suppressing E2F1 activated NPRL2 expression. Cell Biol Int 45, 2510–2520

57. Hollern, D. P., Swiatnicki, M. R., Rennhack, J. P., Misek, S. A., Matson, B. C., McAuliff, A., Gallo, K. A., Caron, K. M., and Andrechek, E. R. (2019) E2F1 Drives Breast Cancer Metastasis by Regulating the Target Gene FGF13 and Altering Cell Migration. Sci Rep 9, 10718

58. Liang, Y. X., Lu, J. M., Mo, R. J., He, H. C., Xie, J., Jiang, F. N., Lin, Z. Y., Chen, Y. R., Wu, Y. D., Luo, H. W., Luo, Z., and Zhong, W. D. (2016) E2F1 promotes tumor cell invasion and migration through regulating CD147 in prostate cancer. Int J Oncol 48, 1650–1658

59. Phillips, A. C., Ernst, M. K., Bates, S., Rice, N. R., and Vousden, K. H. (1999) E2F-1 potentiates cell death by blocking antiapoptotic signaling pathways. Mol Cell 4, 771–781

60. Putzer, B. M., and Engelmann, D. (2013) E2F1 apoptosis counterattacked: evil strikes back. Trends Mol Med 19, 89–98

